# The p97-UBXD8 complex maintains peroxisome abundance by suppressing pexophagy

**DOI:** 10.1101/2024.09.24.614749

**Authors:** Iris D. Montes, Suganthan Amirthagunanathan, Amit S. Joshi, Malavika Raman

**Affiliations:** Department of Developmental Molecular and Chemical Biology, Tufts University School of Medicine, Boston MA; Department of Biochemistry & Cell and Molecular Biology, University of Tennessee, Knoxville, TN

## Abstract

Peroxisomes are vital organelles involved in key metabolic functions in eukaryotic cells. Their significance is highlighted by peroxisome biogenesis disorders; severe childhood diseases marked by disrupted lipid metabolism. One mechanism regulating peroxisome abundance is through selective ubiquitylation of peroxisomal membrane proteins that triggers peroxisome degradation via selective autophagy (pexophagy). However, the mechanisms regulating pexophagy remain poorly understood in mammalian cells. Here we show that the evolutionarily conserved AAA-ATPase p97 and its membrane embedded adaptor UBXD8 are essential for maintaining peroxisome abundance. From quantitative proteomic studies we reveal that loss of UBXD8 affects many peroxisomal proteins. We find depletion of UBXD8 results in a loss of peroxisomes in a manner that is independent of the known role of UBXD8 in ER associated degradation (ERAD). Loss of UBXD8 or inhibition of p97 increases peroxisomal turnover through autophagy and can be rescued by depleting key autophagy proteins or overexpressing the deubiquitylating enzyme USP30. Furthermore, we find increased ubiquitylation of the peroxisomal membrane protein PMP70 in cells lacking UBXD8 or p97. Collectively, our findings identify a new role for the p97-UBXD8 complex in regulating peroxisome abundance by suppressing pexophagy.

## Introduction

Peroxisomes are ubiquitous, dynamic organelles with central roles in lipid metabolism (Mast, Rachubinski, & Aitchison, 2020; Schrader, Kamoshita, & Islinger, 2020). These functions include purine catabolism, bile acid and ether phospholipid synthesis, as well as β- and α- oxidation of very long chain fatty acids (VLCFAs) and branched chain fatty acids (BCFA) (Mast et al., 2020; Schrader et al., 2020; Terlecky & Fransen, 2000). Peroxisomes are also essential for detoxification of reactive oxygen species (ROS) and reactive nitrogen species (RNS) (Chen, Chang, Lin, & Yang, 2020; Mast et al., 2020; Wanders, Ferdinandusse, Brites, & Kemp, 2010). Peroxisome homeostasis is maintained by a group of peroxins (PEX) proteins that coordinate peroxisome biogenesis, import and fission. Given their central importance, loss of peroxisomes or defects in peroxins (PEX) can result in dramatic alterations to the cellular lipidome and can lead to a host of human diseases broadly termed peroxisome biogenesis disorders (PBD) (Aubourg & Wanders, 2013; Mast et al., 2020; Schrader et al., 2020). These include neonatal adrenoleukodystrophy, Zellweger spectrum disorders, and Refsum disease among others, which affect multiple organs with a prominent neurological phenotype (Buchberger, Howard, Proctor, & Bycroft, 2001; Mast et al., 2020; Schrader et al., 2020).

Peroxisome abundance is maintained through de novo biogenesis, fission of mature peroxisomes and degradation through a selective form of autophagy known as pexophagy (Terlecky & Fransen, 2000). Pexophagy can be triggered by environmental stressors, including nutrient deprivation, hypoxia, and high ROS levels (Wei et al., 2021). Several findings suggest that peroxins involved in peroxisome biogenesis as well as the import of matrix proteins contribute to pexophagy (Eun et al., 2018; Wei et al., 2021). For example, overexpression of the peroxisome biogenesis protein PEX3 induces ubiquitylation of peroxisome membrane proteins, leading to pexophagy (Wei et al., 2021; Yamashita, Abe, Tatemichi, & Fujiki, 2014). The peroxisome resident E3 ligase PEX2 can ubiquitylate the import receptor PEX5 as well as the peroxisomal membrane protein PMP70 / ABCD3 during amino acid starvation (Eun et al., 2018; Sargent et al., 2016). Ubiquitylated PEX5 recruits the autophagy adaptor proteins, sequestosome-1 (SQSTM1/p62) and neighbor of BRCA1 gene 1 (NBR1), which target peroxisomes for pexophagy (Léon, Goodman, & Subramani, 2006; Riccio et al., 2019; Sargent et al., 2016; J. Zhang et al., 2015). Despite the importance of maintaining peroxisome homeostasis, the molecular mechanisms regulating pexophagy in mammalian cells are not comprehensively understood (Zalckvar & Schuldiner, 2022).

p97 is an evolutionarily conserved, ATP driven, homohexameric chaperone important for ubiquitin-dependent protein quality control (Neuber, Jarosch, Volkwein, Walter, & Sommer, 2005; Stach & Freemont, 2017). Consecutive ATP hydrolysis by p97 monomers in the hexamer causes unfolding of bound ubiquitylated substrates as they pass through the central pore of the homo-hexamer. While p97 has well established roles in the degradation of ubiquitylated proteins by the 26S proteasome, recent studies indicate that it also participates in early and late steps in autophagy (Ahlstedt, Ganji, & Raman, 2022; Papadopoulos et al., 2017; Tanaka et al., 2010; Zheng, Cao, et al., 2022). p97 associates with over 30 adaptors that interact with its N- or C-termini enabling recruitment of p97 to various organelles and ubiquitylated substrates (Stach & Freemont, 2017). One such adaptor UBXD8 is localized to the ER via a hydrophobic hairpin motif that inserts into the outer leaflet of the ER membrane. At the ER the p97-UBXD8 complex has important functions in ER-associated degradation (ERAD), wherein misfolded proteins in the ER lumen or membrane are ubiquitylated, retro-translocated to the cytosol and degraded by cytosolic proteasomes (Ruggiano, Foresti, & Carvalho, 2014). UBXD8 recognizes ubiquitylated substrates and p97 through its ubiquitin associated (UBA) and ubiquitin X (UBX) domains respectively (Buchberger et al., 2001; Budhidarmo & Day, 2014; H. Kim et al., 2013; Neuber et al., 2005; Schuberth & Buchberger, 2008). Recent studies suggest that endogenous UBXD8 also localizes to and regulates the functions of lipid droplets and mitochondria (Olzmann, Richter, & Kopito, 2013; Song, Herrmann, & Becker, 2021; Zheng, Cao, Yang, & Jiang, 2022).

In previous published work from our lab, we used quantitative tandem mass tag (TMT) proteomics to determine how the cellular proteome is remodeled in UBXD8 knockout (KO) cells generated by CRISPR editing (Ganji et al., 2023). Further analysis of the proteomic dataset led to the surprising finding that numerous peroxisomal proteins were depleted in UBXD8 KO cell lines in comparison to wildtype cells. We explored this unexpected finding further and found that loss of UBXD8 leads to significant reduction in peroxisomes in multiple cell lines. We further show that loss of peroxisomes in UBXD8 KO cells causes an increase in VLCFAs and decreased catalase activity. Moreover, we find endogenous UBXD8 localizes to peroxisomes and loss of p97-UBXD8 increases the degradation of peroxisomes in a manner that is dependent on autophagy and ubiquitylation of peroxisomal membrane proteins. Taken together, we show that the p97-UBXD8 complex controls peroxisome abundance and function by suppressing their degradation through autophagy.

## Results

### Quantitative proteomics identifies altered levels of peroxisomal proteins in UBXD8 knockout cells

In previous published work from our group, we evaluated how the cellular proteome was impacted by loss of UBXD8 (Ganji et al., 2023). We generated UBXD8 KO HeLa and HEK 293T cell lines and performed multiplexed, quantitative proteomics using tandem mass tags (TMT) on triplicate wildtype and UBXD8 KO cells. Details on the analysis can be found in the original manuscript (Ganji et al., 2023). This study enabled the elucidation of a novel role for the p97-UBXD8 complex in regulating ER-mitochondria contact sites by modulating lipid desaturation and membrane fluidity.

Intriguingly, further analysis of this dataset found that numerous peroxisomal proteins were reproducibly depleted in both HeLa and HEK 293T UBXD8 KO cells (61 in HeLa and 65 in HEK 293T) (Figure 1A, Supplementary Figure 1A, B). Ten of these proteins displayed a statistically significant depletion in UBXD8 KO HeLa cells relative to wildtype (log_2_ fold change (wildtype:KO) > 0.65 and −log_10_ p value >1.5) (Figure 1A). Depleted proteins were involved in a variety of peroxisomal processes ranging from import to metabolic functions suggesting global alterations in peroxisomes (Supplementary Figure 1C) (Bagattin, Hugendubler, & Mueller, 2010). We found proteins crucial for biogenesis (PEX3, PEX16 and PEX26) as well as enzymes involved in distinct lipid metabolic reactions such as ATP binding cassette subfamily D member 1 (ABCD1), ATP binding cassette subfamily D member 3 (ABCD3), acyl-CoA dehydrogenase family member 11 (ACAD11) and acyl-CoA oxidase 3 (ACOX3) to be impacted (Figure 1B). We validated our results by immunoblotting for a subset of these peroxisomal proteins in both Hela and HEK293T cells and found significant depletion in both cell lines (Figure 1C-D). Loss of peroxisomal proteins was not due to deficits in transcription as transcript levels of most peroxisomal mRNAs were generally not altered between wildtype and UBXD8 KO cells (Supplementary Figure 1D). Taken together, our quantitative proteomic studies suggest UBXD8 maintains the abundance of many peroxisomal proteins.

**Figure 1:**
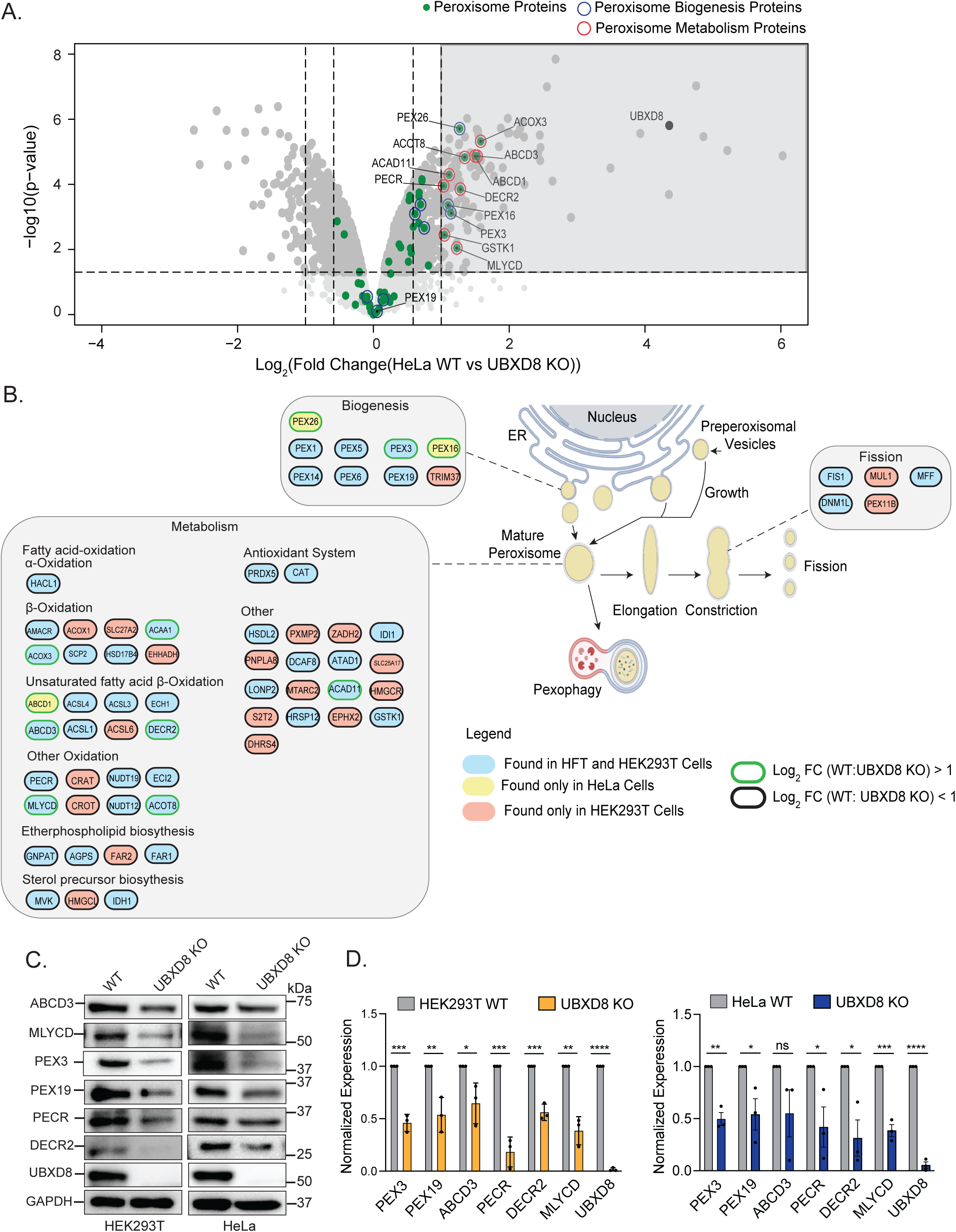
Quantitative proteomics identifies a role for UBXD8 in regulating peroxisome protein abundance. **A.** Volcano plot of the (−log10-transformed *P* value *versus* the log2-transformed ratio of wildtype/ UBXD8 KO) proteins identified from HeLa cells. *n* = 3 biologically independent samples for each genotype. *P* values were determined by empirical Bayesian statistical methods (two-tailed *t* test adjusted for multiple comparisons using Benjamini-Hochberg’s correction method) using the *LIMMA* R package; for parameters, individual *P* values and *q* values, see Supplementary Dataset. Peroxisomal proteins important for biogenesis (dark blue) and metabolism (red) are highlighted. This dataset has been previously published in (Ganji et al., 2023) and is reanalyzed here. **B.** Schematic of tandem mass tag (TMT) proteomic hits in distinct peroxisomal pathways. **C.** Peroxisomal proteins identifies in (A) show reduced expression in UBXD8 KO compared to wildtype HeLa and Hek293T cells. **D.** Quantification of (C). **** : P<0.0001, N=3, Unpaired T-test.

### Deletion of UBXD8 decreases peroxisome abundance

We next asked whether loss of peroxisomal proteins in UBXD8 KO cells impacted peroxisome abundance. Wildtype and UBXD8 KO cells were stained with an antibody to catalase, (a peroxisomal matrix marker) and peroxisome membrane protein 70 (PMP70 / ACBD3). Peroxisome number per cell and size was measured using Aggrecount an automated image analysis script developed in the lab (Klickstein, Mukkavalli, & Raman, 2020). We found that the number of peroxisomes was decreased, and the size of peroxisomes was increased in the absence of UBXD8 in Hela cells (Figure 2A-B, Supplementary Figure 2A-B). Reduction in the number of peroxisomes was further validated in HEK293T UBXD8 KO cells (Supplementary Figure 2D-G), as well as by acute depletion with two distinct siRNAs (Supplementary Figure 2H-J). Notably, ∼10% of UBXD8 KO cells were completely devoid of peroxisomes (Figure 2C, Supplementary Figure 2C). We note that UBXD8 deletion or depletion in other cell lines consistently reduced peroxisome abundance but the increase in peroxisome size was unique to HeLa cells. We used both knockout cells and siRNA depletion in key following studies.

**Figure 2:**
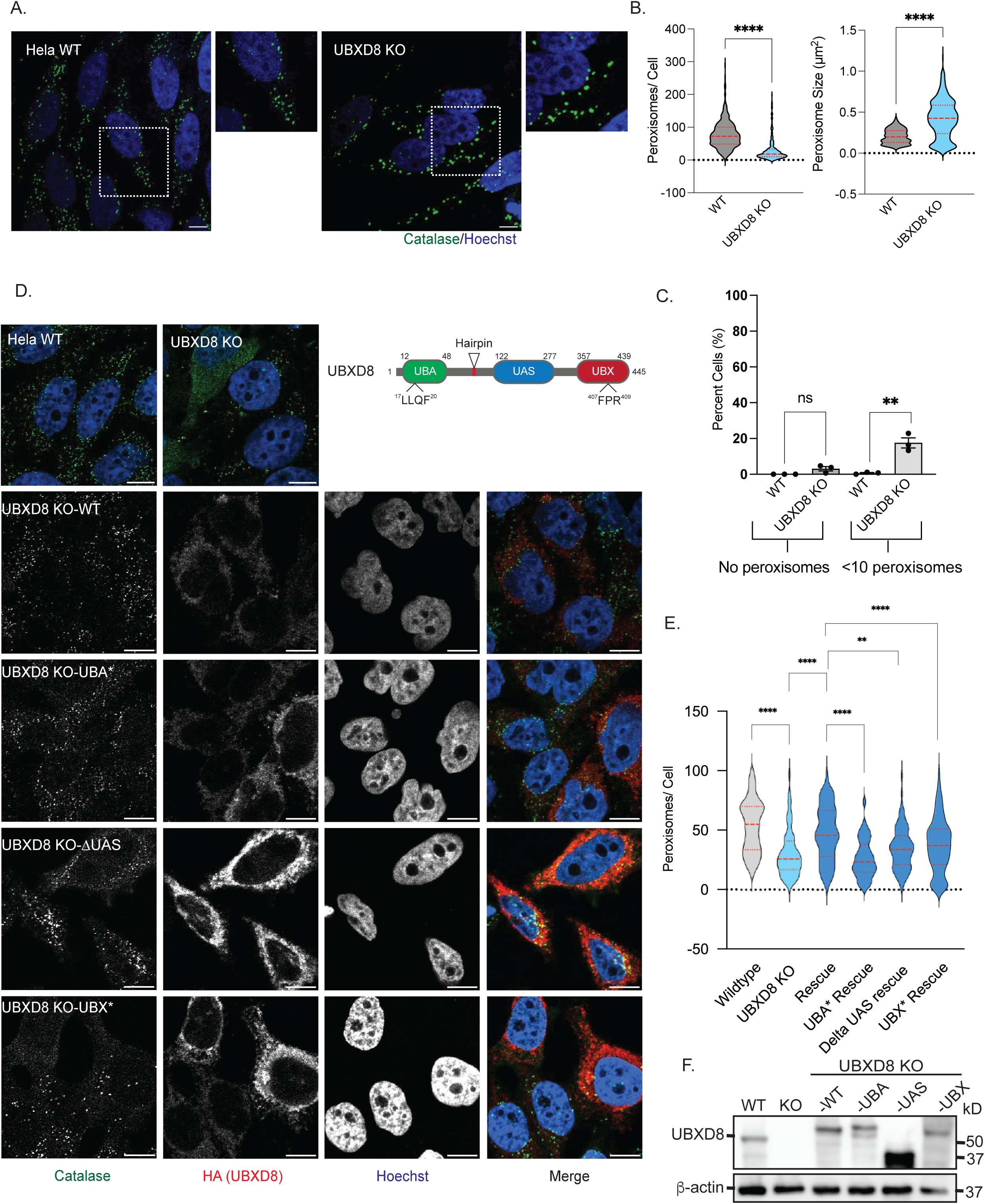
Deletion of UBXD8 perturbs peroxisome abundance. **A.** HeLa wildtype and UBXD8 KO cells stained for peroxisomes using peroxisomal matrix marker catalase **B.** Quantification of average peroxisome per cell and average peroxisome size from (A). At least 100 cells were analyzed in N=3 independent experiments. Violin plot shows median and 95% confidence intervals **** : P<0.0001, N=3, Unpaired T-test. **C.** Peroxisome abundance in wildtype and UBXD8 KO cells that have either no peroxisomes or less then 10 peroxisomes per cell. **D.** Rescue of peroxisome number in UBXD8 KO cells transfected with either UBXD8-HA, UBXD8-UBA*-HA, UBXD8-ΛUAS-HA or UBXD8-UBX*-HA. Cells were stained for peroxisomes using peroxisomal matrix marker catalase. **E.** Quantification of average peroxisome per cell from HeLa wildtype and UBXD8 KO cells as well as UBXD8 KO cells transfected with either UBXD8-HA, UBXD8-UBA*-HA, UBXD8-ΛUAS-HA or UBXD8-UBX*-HA (D). Peroxisome numbers were quantified in cells expressing HA-tagged UBXD8 constructs only. At least 100 cells were analyzed in N=3 independent experiments. Violin plot shows median and 95% confidence intervals **, **** : P<0.05, 0.0001. Two-way ANOVA with Tukey’s multiple comparisons test. **F.** Immunoblots of constructs transfected. Scale bar is 10μM.

In addition to its UBA and UBX domains, UBXD8 also contains a central UAS domain that has been reported to interact with long chain unsaturated fatty acids (H. Kim et al., 2013). We next asked if these domains were important in the role of UBXD8 in regulating peroxisome abundance. UBXD8 KO cells were transfected with HA-tagged wildtype UBXD8 or individual UBA or UBX domain point mutants that we have previously verified to lose interaction with ubiquitin and p97 respectively (Ganji et al., 2023). We also transfected cells with a UAS domain deletion mutant. While expression of wildtype UBXD8 rescued peroxisome abundance in UBXD8 KO cells (Figure 2 D-F), point mutants in the UBA or UBX domain or deletion of the UAS domain did not rescue as well as wildtype (Figure 2D-F). In summary, we find that UBXD8 deletion decreases peroxisome abundance in multiple cell lines in a manner dependent on ubiquitin interaction and p97 recruitment.

### Loss of peroxisomes in UBXD8 KO cells is not due to ER stress or ERAD inhibition

UBXD8 has important functions in ERAD to help prevent or alleviate ER stress (Schuberth & Buchberger, 2008). Given that the ER can regulate peroxisome biogenesis, it is possible that loss of UBXD8 and subsequent inhibition of ERAD could cause ER stress thus leading to perturbed peroxisome abundance. We therefore systematically tested if loss of peroxisomes in UBXD8 KO cells was a secondary consequence of ER stress. First, we asked if depletion of two major E3 ligases that execute ERAD; HMG CoA reductase degradation 1 (HRD1) and GP78 (Schulz et al., 2017; Zhang, Xu, Liu, & Ye, 2015) impacted peroxisome abundance. HRD1 depletion with two siRNAs had small but significant changes in peroxisome numbers but there was no concordance in phenotype between the two siRNAs and it did not phenocopy the UBXD8 deletion phenotype (Figure 3A and Supplementary Figure 3A-B). Similarly, a CRISPR generated GP78 KO HEK293T cell line (Bersuker et al., 2018) had a small increase in peroxisome abundance (Figure 3B and Supplementary Figure 3C-D). Thus, loss of ERAD E3s did not recapitulate the UBXD8 KO phenotype. Second, we asked if alternate p97 ERAD adaptors such as UBXD2 (Liang et al., 2006) regulated peroxisome abundance. We measured peroxisome number and size in a HeLa UBXD2 KO cell line and found no significant change in peroxisome abundance (Figure 3C-D). Third, we asked if simply increasing ER stress impacted peroxisome numbers. We treated cells with tunicamycin which causes protein misfolding in the ER by preventing N-linked glycosylation but observed no impact on peroxisome number (Supplementary Figure 3E-G). Finally, we asked if there was increased ER stress in UBXD8 KO cells by assessing levels of the ER chaperone binding immunoglobulin protein (BiP), activating transcription factor 4 (ATF4) by immunoblot, and transcript levels of *x box binding protein 1 spliced* (*xbp1_s_*) by quantitative real time PCR. Wildtype cells were treated with dithiothreitol (DTT), to induce ER stress as a positive control. We found an increase in BiP, ATF4, and *xbp1_s_* levels in wildtype cells treated with DTT. However, the levels of these markers were unchanged in UBXD8 KO cells (Figure 3E-F). Thus, we conclude that the decrease in peroxisome abundance in UBXD8 KO cells is not due to altered ER protein homeostasis.

**Figure 3:**
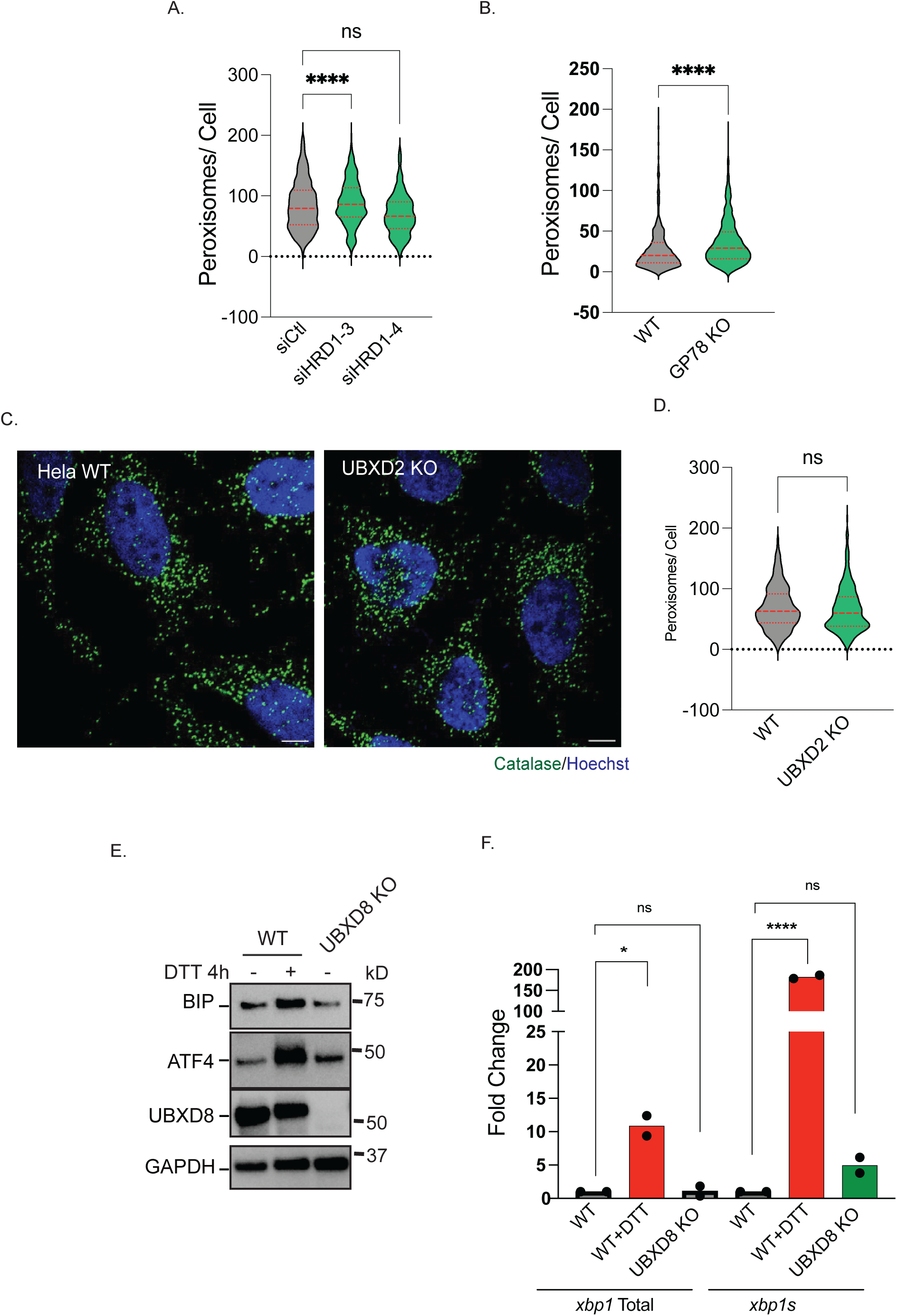
ER stress does not contribute to loss of peroxisomes in UBXD8 null cells. **A.** Hela cells were transfected with siRNAs to HRD1, and cells were stained for catalase (see also Supplementary Figure 3). Quantification of number of peroxisomes per cell, at least 100 cells were analyzed in N=3 independent experiments. Violin plot shows median and 95% confidence intervals. ns: not significant, **** : P<0.0001. One-way ANOVA with Dunnett’s multiple comparisons test. **B.** Quantification of peroxisome per cell in HEK293T wildtype and GP78 KO cells. At least 100 cells were analyzed in N=3 independent experiments. Violin plot shows median and 95% confidence intervals. **** : P<0.0001. Unpaired T-test. **C.** HeLa wildtype and UBXD2 KO cells stained for peroxisomes using catalase. **D.** Quantification of peroxisome per cell from (C). At least 100 cells were analyzed in N=3 independent experiments. Violin plot shows median and 95% confidence intervals. NS: not significant. Unpaired T-test. **E.** Immunoblots of Hela wildtype (untreated or treated with DTT) and UBXD8 KO for ER stress markers BiP and ATF4. N=3, **F.** rt-qPCR of *xbp1* total and *xbp1 spliced (xbp1s)* mRNA transcripts in wildtype and UBXD8 KO cells treated with 1.5 mM DTT for 4 hours. N=3. NS: Not significant, *, **** : P < 0.01, 0.0001. Two-way ANOVA with Dunnett’s post-hoc analysis. Scale bar is 10μM.

### UBXD8 KO cells have dysfunctional peroxisomes

Loss of peroxisomes (for example in PBD) causes decreased plasmalogen and cholesterol levels as well as an accumulation of VLCFAs and phytanic acid (Aubourg & Wanders, 2013; Faust & Kovacs, 2014). We had previously performed lipidomics in wildtype and UBXD8 KO cells and re-analyzed that dataset to evaluate acyl chain lengths in the major classes of lipids. (Ganji et al., 2023). We found a significant accumulation of very long chain fatty acyl chains in cholesterol esters (CE), triacyglycerides (TG) and distinct phospholipid species in UBXD8 KO cells compared to wildtype cells (Figure 4A-B). These lipids were conjugated with acyl chains ranging from 28 to 56 hydrocarbons indicative of an increase in very long chain fatty acids (Figure 4A-B). We also investigated peroxisome function by evaluating catalase activity in wildtype and UBXD8 KO cells. We found a decrease in catalase activity in UBXD8 KO cells compared to wildtype cells (Figure 4D). However, catalase levels were comparable between wildtype and UBXD8 KO cells (Figure 4E). This is likely due to the known redistribution of catalase to the cytoplasm in cells lacking peroxisomes. Thus, the loss of peroxisomes in UBXD8 KO cells leads to perturbed lipid metabolism and decreased catalase activity.

**Figure 4:**
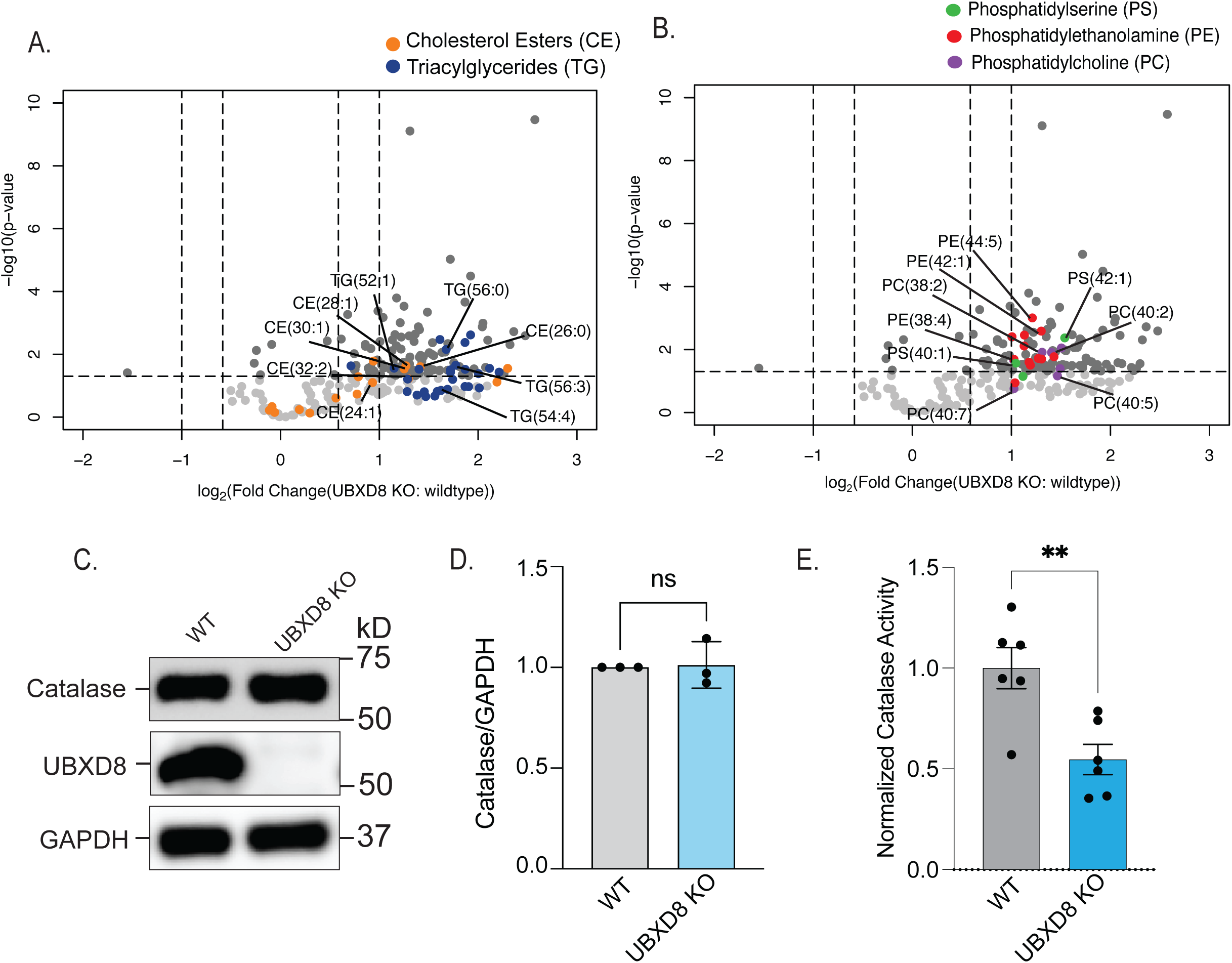
Depletion of UBXD8 results in increase of VLCFAs and a loss in catalase activity. **A.** Volcano plot of the total cholesterol esters and triacylglycerol species identified using lipidomics of whole cell extracts of HEK-293T cells (−log10-transformed P value versus the log2-transformed ratio of UBXD8 KO: wildtype). VLCFA species indicated for CE (orange) and TG (dark blue). This dataset has been previously published in (Ganji et al., 2023) and is reanalyzed here. **B.** VLCFA species indicated for phosphatidylserine (PS) (green), phosphatidylethanolamine (PE) (red) and phosphatidylcholine (PC) (violet). Lipids were measured by LC-MS/MS following normalization by total protein amount. (n ≥ 3 biologically independent experiments were performed, each with duplicate samples). This dataset has been previously published in (Ganji et al., 2023) and is reanalyzed here. **C.** Immunoblots of catalase levels in whole cell lysates of HeLa wildtype and UBXD8 KO cells. **D.** Quantification of catalase levels in (C). N=3 independent experiments. NS: not significant. Unpaired T-test. **E.** Catalase activity was quantified using a commercial kit. N=3 independent experiments. ** : P<0.0001, Unpaired T-Test.

### UBXD8 localizes to peroxisomes

While UBXD8 is localized to the ER, recent studies have found that endogenous UBXD8 can also localize to mitochondria and lipid droplets to regulate organelle function (Zheng, Cao, et al., 2022). Intriguingly, a recent proteomics analysis of purified peroxisomes identified UBXD8 as a putative peroxisome localized protein (Ray et al., 2020). We therefore asked whether UBXD8 could localize to peroxisomes. We transiently transfected COS-7 and HeLa cells with UBXD8-GFP and RFP-SKL (a type 1 peroxisome targeting sequence). Cells were also labelled with BODIPY (665/676) to label lipid droplets. As previously reported, GFP-UBXD8 was found on the surface of lipid droplets (Figure 5A). Notably, we observed robust localization of UBXD8 to peroxisomes, with UBXD8 forming a ring around peroxisome labelled with RFP-SKL (Figure 5A). We also found endogenous UBXD8 localized to peroxisomes (Supplementary Figure 4A). To characterize which UBXD8 domains contribute to peroxisome localization, we transfected UBXD8 KO cells with wildtype or UBXD8 mutants in the UBA, UAS and UBX domains (both point and deletion mutants). UBXD8 without the UBA domain less efficiently targeted to peroxisomes (Figure 5B-C and Supplementary Figure 4 B-C). These studies suggest that UBXD8 localizes to peroxisomes in a manner dependent on ubiquitin association.

**Figure 5:**
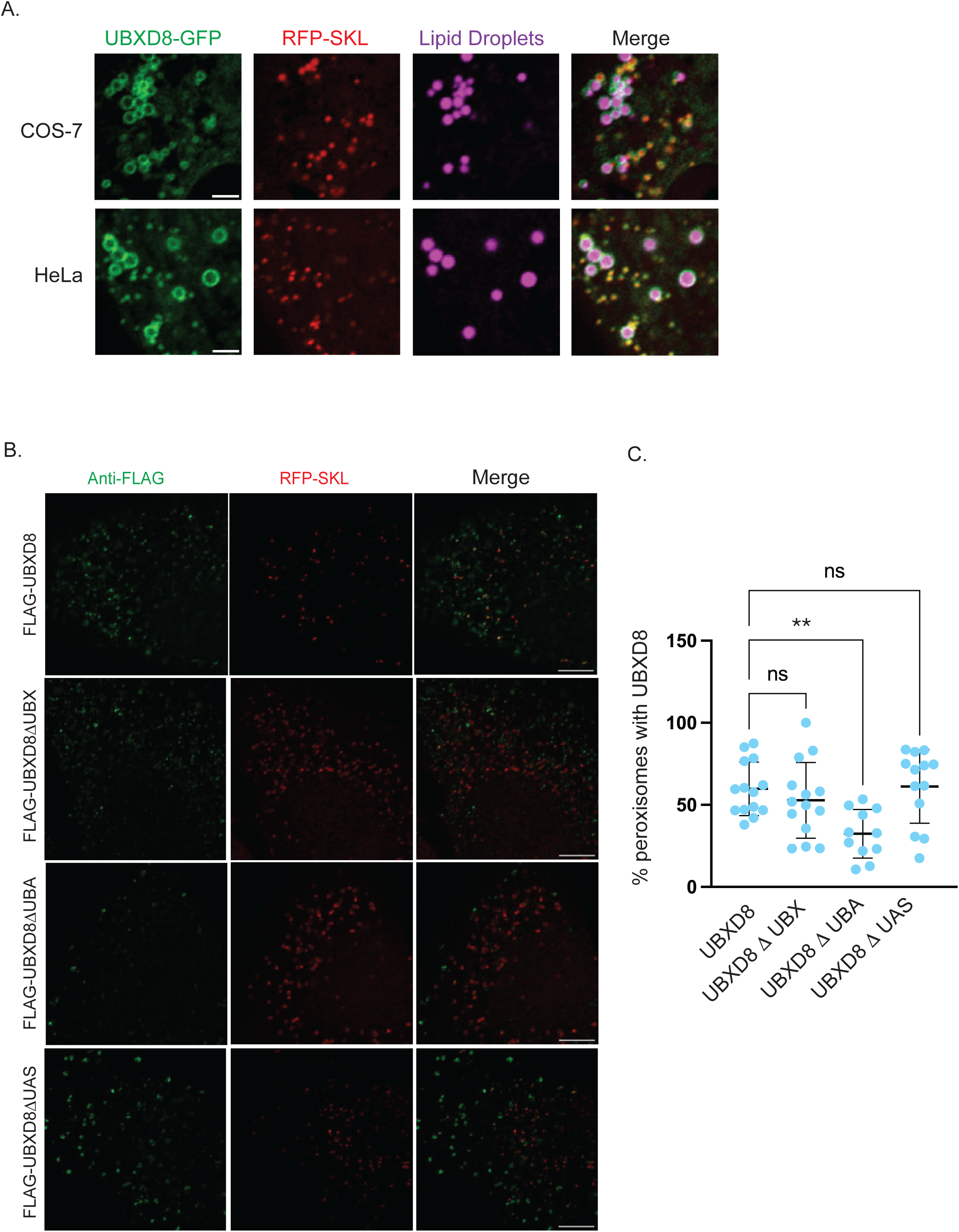
UBXD8 localizes to peroxisomes. **A.** GFP-UBXD8 and RFP-SKL were transiently transfected into COS-7 or HeLa cells and stained with BODIPY (665/676) to label lipid droplets. **B.** HeLa cells were transfected with FLAG-tagged wildtype UBXD8 or UBXD8 domain deletions (UBA, UAS and UBX) (in green) and RFP-SKL (in red). **C**. Quantification of (B) showing number of peroxisomes with UBXD8 localization. 15-20 cells were analyzed in N=3 independent experiments. Scatter plot shows mean and std.dev. ns: not significant, ** : P<0.001. One-way ANOVA with Dunnett’s multiple comparisons test. Scale bar is 5μM.

### p97-UBXD8 suppress pexophagy by targeting ubiquitylated PMP70

Given the role of p97-UBXD8 in the extraction and degradation of organelle localized membrane proteins and the loss of peroxisomes in UBXD8 KO cells, we asked if this complex participated in pexophagy. We used a peroxisomal flux reporter that consisted of a tandem chimera of mCherry and eGFP fused to the peroxisome membrane targeting sequence of PEX26 (J. Zhang et al., 2015; Zheng, Chen, Liu, Zhong, & Zhuang, 2022). Both mCherry and eGFP fluorescence (yellow puncta) is observed for healthy peroxisomes or those residing in autophagosomes. However, peroxisomes in lysosomes only harbor the mCherry signal as the GFP fluorophore is quenched in the acidic lumen of the lysosome (Figure 6A). Wildtype and UBXD8 KO cells were transiently transfected with eGFP–mCherry–PEX26 and cells were treated with Torin1 a pan-mTOR inhibitor (J. Zhang et al., 2015; Zheng, Chen, et al., 2022) to stimulate pexophagy. As expected Torin1 treatment resulted in activation of pexophagy and loss of peroxisomes in wildtype cells and UBXD8 KO cells (Supplementary Figure 5A-B). We quantified the ratio of GFP:mCherry and found a significant loss in eGFP signal in UBXD8 KO cells compared to wildtype cells under basal conditions which was further stimulated in the presence of Torin1 (Figure 6B-C). To assess a role for p97 in this process, we used siRNA to knockdown p97 and found that loss of p97 also enhanced pexophagy in untreated and Torin1 treated cells (Figure 6D-F). Together our results indicate that both p97 and UBXD8 suppress peroxisome degradation.

**Figure 6:**
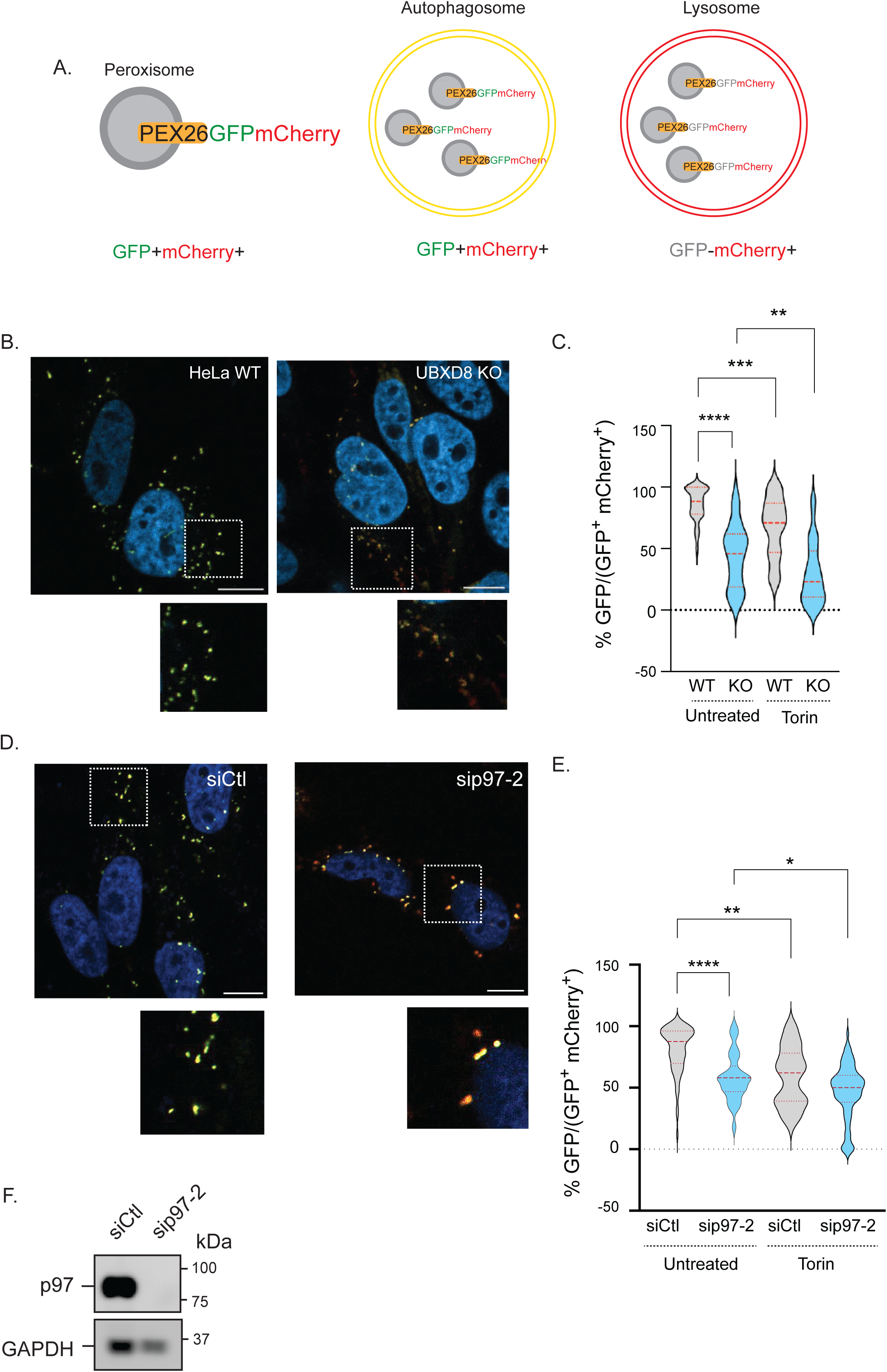
p97-UBXD8 suppress pexophagy. **A.** Schematic for pexophagy flux reporter. **B.** Representative images of wildtype and UBXD8 KO cells transfected with GFP-Cherry-PEX26. **C.** Wildtype and UBXD8 KO cells were transfected with the flux reporter and treated with 150 nM Torin1 for 18 hours. Quantification showing the ratio of GFP: (GFP^+^mCherry^+^) in Hela wildtype and UBXD8 KO cells. 50-100 cells were analyzed in N=3 independent experiments. Violin plot shows median and 95% confidence intervals. **, ***, **** : P<0.01, 0.001, 0.0001. Two-way ANOVA with Šidáks multiple comparisons test. **D.** Representative images of HeLa control or p97 siRNAs and GFP-Cherry-PEX26. **E.** Quantification showing the ratio of GFP: (GFP^+^mCherry^+^) in Hela control, or p97 depleted cells. 50-100 cells were analyzed in N=3 independent experiments. Violin plot shows median and 95% confidence intervals. *, **, **** : P<0.05, 0.001, 0.0001. Two-way ANOVA with Tukey’s multiple comparisons test. **F.** Immunoblot showing p97. Scale bar is 10 μM.

To confirm that increased autophagy was responsible for loss of peroxisomes in UBXD8 loss of function cells, we depleted ATG5, an essential autophagy protein responsible for phagophore elongation (Kuma et al., 2004; Mizushima et al., 2001). We found that depletion of ATG5 in UBXD8 loss of function cells was sufficient to rescue peroxisome abundance (Figure 7A-C and Supplementary Figure 6A-C). To further validate this finding, we over-expressed GFP-USP30, deubiquitylating enzyme necessary for suppressing pexophagy during amino acid starvation (Marcassa et al., 2018; Riccio et al., 2019). UBXD8 KO cells expressing GFP-USP30 had increased peroxisome numbers compared to untransfected cells (Figure 7D-F). NBR1 is the key autophagy receptor for pexophagy. We immunostained wildtype and UBXD8 KO cells with for NBR1 and catalase and found increased co-localization of peroxisomes with NBR1 in UBXD8 KO cells (Figure 7G-H).

**Figure 7:**
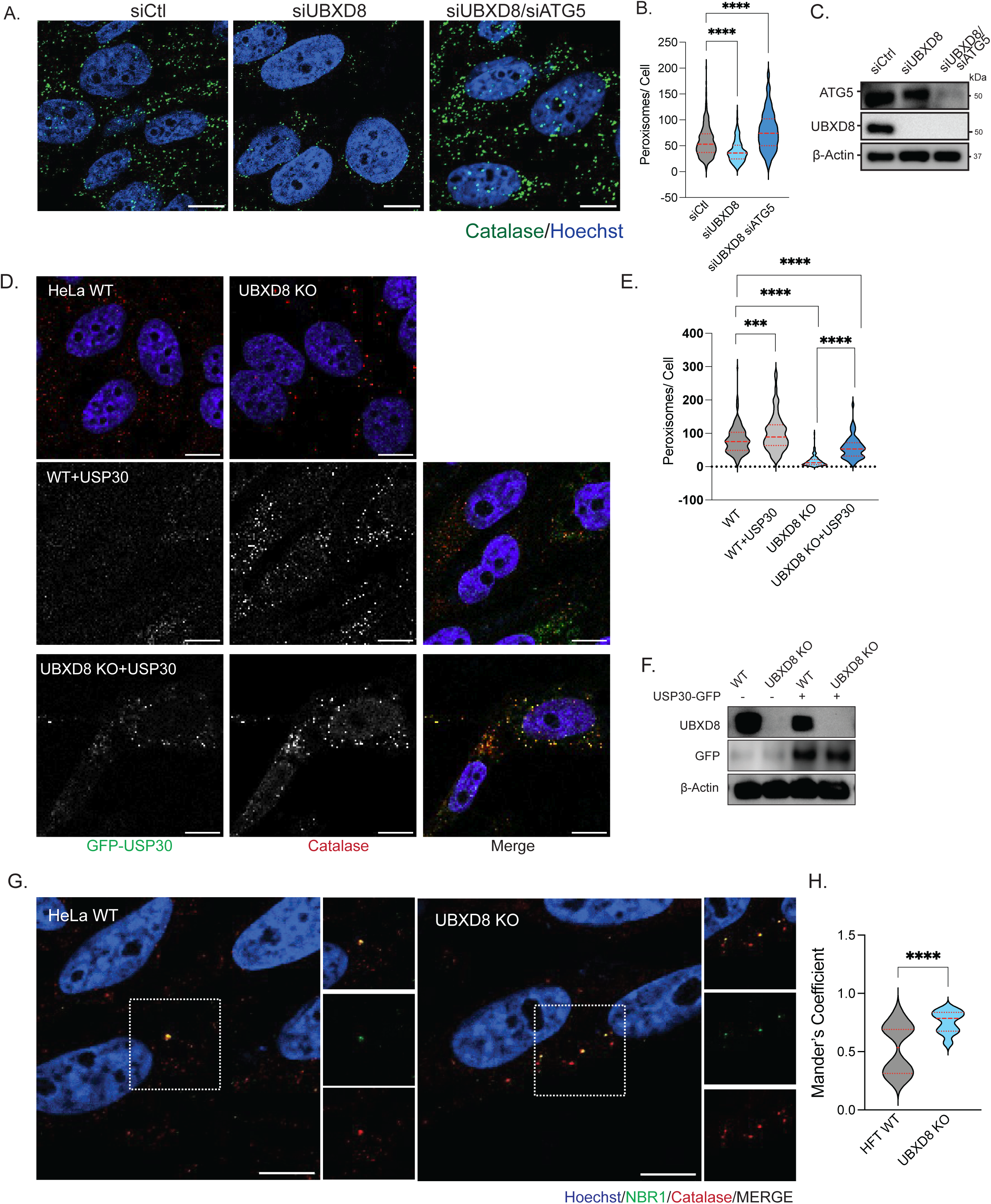
Depletion of ATG5 and over-expression of USP30 rescues pexophagy in UBXD8 deleted cells. **A.** Representative images of HeLa cells transfected with control and UBXD8 or ATG5 siRNAs and stained for catalase. **B.** Quantification of peroxisomes per cell from (A). 50-100 cells were analyzed in N=3 independent experiments. Violin plot shows median and 95% confidence intervals. **** = P<0.0001. Two-way ANOVA with Dunnett’s multiple comparisons test. **C.** Immunoblot of UBXD8 and ATG5. **D.** Representative images of HeLa cells (wildtype and UBXD8 KO) transfected with GFP-USP30. Cells were stained for catalase. **E.** Quantification of peroxisomes per cell in GFP-USP30 transfected cells. 50-100 cells were analyzed in N=3 independent experiments. Violin plot shows median and 95% confidence intervals. ***, **** = P< 0.001, 0.0001. Two-way ANOVA with Dunnett’s multiple comparisons test N=3, 2-way ANOVA. **F.** Immunoblot of GFP-USP30 expression. **G.** Wildtype and UBXD8 KO HeLa cells were stained with NBR1 and catalase. **H.** Mander’s colocalization of images in (G). 100-120 cells were analyzed in N=3 independent experiments. Violin plot shows median and 95% confidence intervals. **** :P< 0.0001. Students unpaired T-test. **F.** Immunoblot of GFP-USP30 expression. Scale bar is 10 μM (A and D) and 5 μM (G).

Ubiquitylation of peroxisomal membrane proteins is the signal for pexophagy. A number of studies have found that the membrane protein PMP70 and the import receptor PEX5 are ubiquitylated under various settings to stimulate pexophagy (Ott et al., 2022; Sargent et al., 2016; J. Zhang et al., 2015; Zheng, Chen, et al., 2022). Indeed, we found that treatment of cells with Torin-1 decreased the half-life of PMP70 whereas the autophagy inhibitor Bafilomycin A prolonged it (Figure 8A-B). Moreover, we found increased ubiquitylation of PMP70 in cells treated with the proteasome inhibitor Bortezomib or Bafilomycin A under conditions that stimulated pexophagy (Figure 8C). We therefore asked whether ubiquitylation of PMP70 was perturbed in cells depleted for p97 or UBXD8. We find that depletion of either p97 or UBXD8 resulted in an increase in ubiquitylated PMP70 (Figure 8D). Collectively, our studies suggest that loss of p97-UBXD8 causes failure to degrade PMP70 leading to the loss of peroxisomes due to increased pexophagy (Figure 8E).

**Figure 8:**
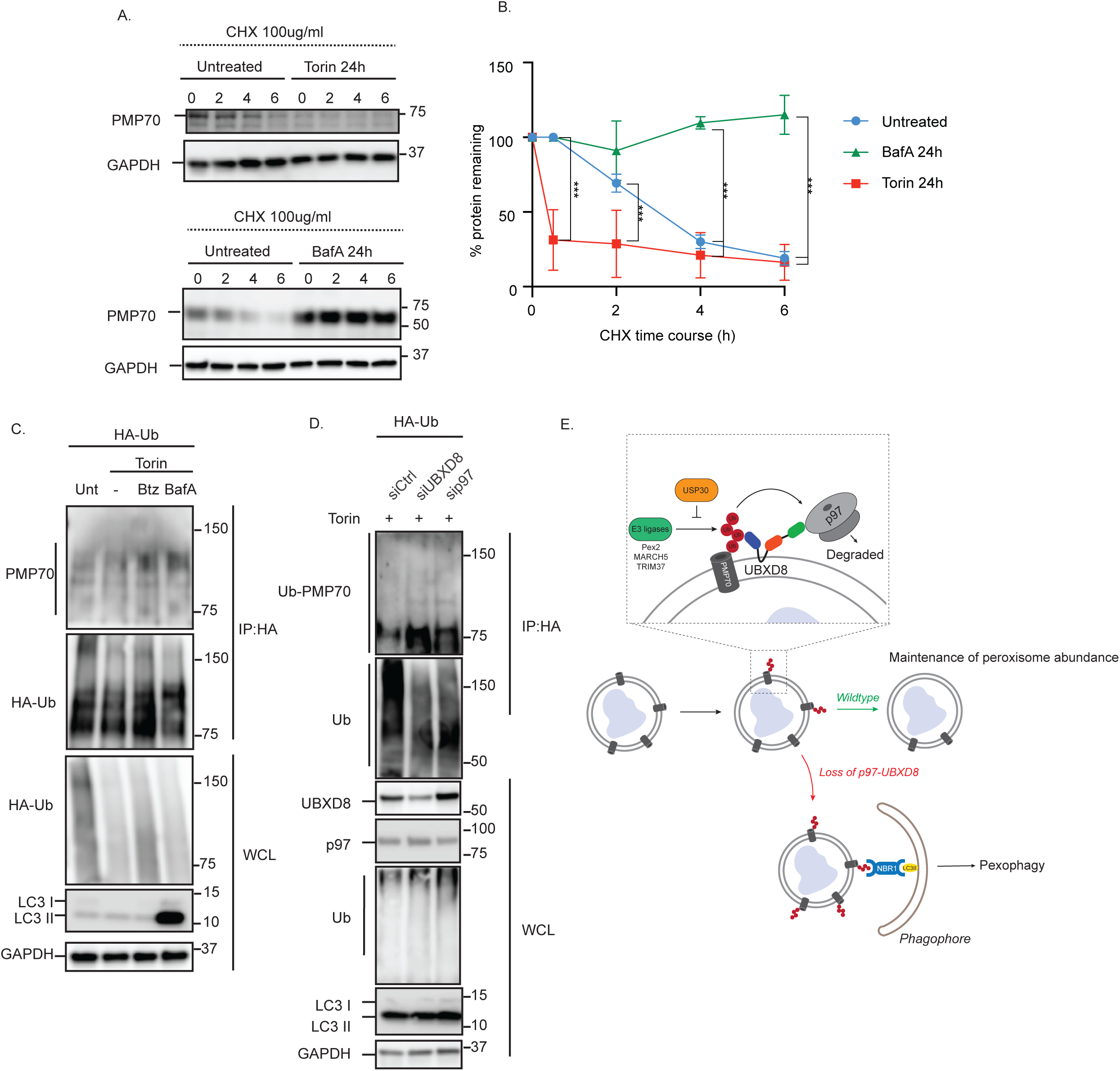
Persistent PMP70 ubiquitylation in cells depleted of p97-UBXD8. **A.** HEK293T cells were treated with 150 nM Torin or 50 nM bafilomycin A (BafA) for 24 hours and then with 100 μg/ ml cycloheximide for the indicated time points to stop translation. Immunoblots showing PMP70 half-life. **B.** Quantification of PMP70 levels normalized to GAPDH from (A). N=3 independent experiments. **, *** : P< 0.01, 0.001. Two-way ANOVA with Tukey’s multiple comparisons test. **C.** HEK293T cells were transfected with HA-ubiquitin and treated with 150 nM Torin-1 in the presence or absence of 5 μM Bortezomib (Btz) or 50 nM BafA for 24 hours. Cells were lysed in SDS, and denaturing HA immunoprecipitations were performed. Immunoblots of indicated proteins showing increased ubiquitylation of PMP70 in cells treated with Btz or BafA. N=3 independent experiments. **D.** HEK293T cells were transfected with HA-ubiquitin and siRNAs to control, UBXD8 or p97. Cells were treated with 150 nM Torin-1 for 24 hours. Cells were lysed in SDS, and denaturing HA immunoprecipitations were performed. Immunoblots of indicated proteins showing increased ubiquitylation of PMP70 in cells depleted of UBXD8 and p97. N=3 independent experiments. E. Model showing p97-UBXD8 suppression of pexophagy.

## Discussion

In this study we find a novel role for p97 and its adaptor UBXD8 in maintenance of peroxisome numbers by suppressing pexophagy. Our previous study examining how the proteome was altered by deletion of UBXD8 identified widespread loss of peroxisomal proteins in UBXD8 KO cells (Figure 1). We show that the loss of peroxisomal proteins is due to the significant depletion of peroxisomes (Figure 2). UBXD8 has been extensively characterized as a p97 adaptor in ER-associated degradation (ERAD). The ER is essential for peroxisome homeostasis; and acts as the site for biogenesis of pre-peroxisomal vesicles (P. K. Kim, Mullen, Schumann, & Lippincott-Schwartz, 2006; van der Zand, Gent, Braakman, & Tabak, 2012) and a donor of membrane lipids via ER-peroxisome contact sites (Joshi, 2021; Kors et al., 2022). Thus, it is possible that peroxisome loss in UBXD8 deleted cells was a secondary consequence of perturbed ER function. However, using a suite of complementary studies we conclude that the loss of peroxisomes in UBXD8 deficient cells is independent of its role in ERAD (Figure 3). Contact sites between the ER and peroxisomes are reported to provide lipids for growth of their peroxisomal membranes (Shai, Schuldiner, & Zalckvar, 2016, Joshi, 2021; Kors et al., 2022). The peroxisomal membrane protein acyl-CoA binding domain containing 5 (ACBD5) tethers peroxisomes to the ER through interaction with vesicle-associated membrane protein-associated proteins B (VAPB) (Kors et al., 2022). This tethering complex may help facilitate lipid transport from the ER to peroxisomes. Notably, we have shown that p97-UBXD8 regulates ER-mitochondria contact sites and UBXD8 is enriched at these contacts (Ganji et al., 2023). It remains to be determined whether UBXD8 also localizes to and regulates ER-peroxisome contacts and warrants future investigation.

UBXD8 is localized on mitochondria and lipid droplets where it has been demonstrated to recruit p97 for the extraction and degradation of membrane proteins (Olzmann et al., 2013; Zheng, Cao, et al., 2022). This feature is reminiscent of its role in ERAD. In this study we also identified UBXD8 localized to peroxisomes in a manner dependent on ubiquitin association (Figure 5). How UBXD8 localizes to peroxisomes is currently unknown. It may be by directly inserted into the peroxisomal membrane after translation or by migration from the ER. Intriguingly, a previous study by the Kopito group found that Pex19 was essential for inserting UBXD8 into the ER (Schrul & Kopito, 2016). That study also found that sites of insertion were in close apposition to peroxisomes using a semi-permeabilized system and in vitro translated UBXD8. While we observe localization of UBXD8 to peroxisomes, given the resolution limitation of our imaging studies it is possible that these peroxisomes may be in contact ER-localized UBXD8.

Several lines of evidence support the finding that the loss of peroxisomes in UBXD8 null cells is due to increased pexophagy (Figures 6-8). Using a peroxisomal flux reporter we show that loss of p97 or UBXD8 increases flux of peroxisomes through autophagy in untreated cells and increases under conditions that stimulate pexophagy. We further show that peroxisome numbers in UBXD8 null cells can be restored to wildtype levels by depleting the autophagy initiating protein ATG5 or by over-expressing the deubiquitylate USP30. p97 has multiple unique roles in autophagy. p97 associates with the deubiquitylase ataxin-3 to stabilize Bcl-2 interacting protein (BECLIN1), a key constituent of phosphatidylinositol-3-kinase (PI3K) complex I (Riccio et al., 2019). p97 has also been shown to regulate the fusion of autophagosomes with lysosomes, however the mechanism remains poorly understood (Ju et al., 2009). p97 regulates other selective autophagy processes such as mitophagy and lysophagy (Ahlstedt, Ganji, & Raman, 2022; Papadopoulos et al., 2017; Tanaka et al., 2010; Zheng, Cao, et al., 2022), so perhaps it is not surprising that it also regulates the selective degradation of peroxisomes. However, we would like to draw a key distinction for the role of p97 in pexophagy versus other forms of selective autophagy. While loss of p97 activity inhibits mitophagy and lysophagy, we find that p97 depletion *enhances* pexophagy. In mitophagy and lysophagy, p97 mediates the selective extraction of K48-linked ubiquitylated substrates for proteasomal degradation. This degradation may help expose K63-linked ubiquitylated substrates for efficient recruitment of autophagy receptors (Papadopoulos et al., 2017). Notably, a recent study found that another AAA-ATPase ATAD1 may be involved in the degradation of PEX5 when it cannot recycle efficiently (Ott et al., 2022). Thus, multiple mechanisms likely exist to maintain peroxisome abundance and functionality.

Ubiquitylation of peroxisomal proteins serves as a signal for autophagic degradation. Early studies demonstrated that ubiquitin fused to PEX3 or PMP34 on the peroxisomal membrane was sufficient for recognition of peroxisomes by the autophagy receptor p62 and NBR1 (P. K. Kim, Hailey, Mullen, & Lippincott-Schwartz, 2008, Deosaran et al., 2013). Indeed, we find greater co-localization of NBR1 with peroxisomes in UBXD8 null cells (Figure 7). Several E3 ligases have been identified to ubiquitylate peroxisomal proteins. PEX2, a component of the peroxisomal E3 ubiquitin ligase complex ubiquitylates the import receptor PEX5 and the membrane protein PMP70 (Sargent et al., 2016). MARCH5 has also been shown to ubiquitylate PMP70 (Zheng, Chen, et al., 2022). We find that loss of p97 and UBXD8 causes the accumulation of ubiquitin modified PMP70 (Figure 8). PMP70 and PEX5 are the most well characterized peroxisomal proteins whose ubiquitylation triggers pexophagy. Given the promiscuity of other autophagy E3 ubiquitin ligases, PEX2 and other E3 ligases likely ubiquitylate additional peroxisomal membrane proteins that remain to be identified. While a number of studies have systematically identified ubiquitylated proteins on the surface of mitochondria and peroxisomes (Deosaran et al., 2013; Ordureau et al., 2014; Sargent et al., 2016) this has not been done in the case of peroxisomes. It is likely that more peroxisomal membrane proteins are ubiquitylated than currently appreciated. An inventory of these proteins is needed to fully understand the mechanisms regulating pexophagy.

Our findings demonstrate the p97-UBXD8 complex as a novel regulator of peroxisome quality control, highlighting the intricate balance of degradation and recycling necessary to maintain peroxisome homeostasis.

## Figure Legends (Main and supplemental)

**Supplementary Figure 1: Quantitative proteomics of wildtype and UBXD8 KO cells identified loss of peroxisomal proteins. A.** Volcano plot of the (−log10-transformed *P* value *versus* the log2-transformed ratio of wildtype/ UBXD8 KO) proteins identified from HEK293T cells. *n* = 3 biologically independent samples for each genotype. *P* values were determined by empirical Bayesian statistical methods (two-tailed *t* test adjusted for multiple comparisons using Benjamini-Hochberg’s correction method) using the *LIMMA* R package; for parameters, individual *P* values and *q* values, see (Ganji et al., 2023) for dataset. Peroxisomal proteins are shown in green filled circles. Outlines indicate proteins involved in biogenesis (dark blue) and metabolism (red). **B.** Correlation of two HEK293T UBXD8 KO clones used for TMT analysis. **C.** Bubble plot representing significantly enriched GO clusters identified from TMT proteomics of CRISPR UBXD8 KO (black) and wildtype (blue) cells. Size of the circle indicates the number of genes identified in each cluster. **D.** RT-qPCR assessment of different peroxisomal transcripts in wildtype and UBXD8 KO cells. N=3 independent experiments. Graph shows mean and std.dev. NS: Not significant. *, **, ***: P < 0.05, 0.01, 0.001 Students unpaired T-test

**Supplementary Figure 2: UBXD8 deletion leads to loss of peroxisomes in multiple cell lines. A.** HeLa wildtype and UBXD8 KO cells stained for peroxisomes using peroxisomal membrane protein PMP-70. **B.** Quantification of peroxisomes per cell and peroxisome size in (A). 50-100 cells were analyzed in N=3 independent experiments. Violin plot shows median and 95% confidence intervals. **** = P< 0.0001 unpaired T test. **C.** Peroxisome numbers in wildtype and UBXD8 KO cells that have either no peroxisomes or less then 10 peroxisomes per cell. NS: Not significant, * = P< 0.05 unpaired T test. **D.** HEK293T wildtype and UBXD8 KO cells stained for peroxisomes using peroxisomal matrix protein catalase. **E.** Quantification of peroxisomes per cell and peroxisome size from (D). 50-100 cells were analyzed in N=3 independent experiments. Violin plot shows median and 95% confidence intervals. ****: P <0.0001, N=3, Unpaired T-test. **F and G.** Same as D and E but stained for PMP70. **H.** HeLa cells were transfected with control r two different UBXD8 siRNAs and stained for catalase. **J.** Quantification of peroxisomes per cell and peroxisome size from (H). 50-100 cells were analyzed in N=3 independent experiments. Violin plot shows median and 95% confidence intervals. NS: Not significant, ***, **** : P <0.001, 0.0001. Unpaired T-test. **J.** Immunoblot showing UBXD8 depletion. Scale bar is 10 μM (A, D, F) and 5 μM (H).

**Supplementary Figure 3: ER stress does not contribute to loss of peroxisomes in UBXD8 KO cells. A.** HeLa cells were transfected with control or two different HRD1 siRNAs and stained for catalase. **B.** Immunoblot showing depletion of Hrd1. **C.** HEK293T wildtype cells and GP78 KO cells stained for catalase. **D.** Immunoblot of GP78 KO. **E.** HFT wildtype cells treated with tunicamycin (2.5μM for 4hrs) and stained for catalase. **F.** Quantification of peroxisomes per cell from (E). 50-100 cells were analyzed in N=4 independent experiments. Violin plot shows median and 95% confidence intervals. NS: Not significant. Unpaired T-test. **G.** Immunoblot of BiP induction in tunicamycin (Tu) treated cells. Scale bar is 10 μM.

**Supplementary Figure 4: Endogenous UBXD8 localizes to peroxisomes in a ubiquitin dependent manner**. **A.** HeLa cells were transiently transfected with SKL-GFP to label peroxisomes and immunostained with an antibody to UBXD8 (red). **B.** HeLa cells were transiently transfected with FLAG-tagged wildtype UBXD8 or UBXD8 domain mutants (UBA and UBX) (in green) and RFP-SKL (in red). **C**. Quantification of (B) showing number or peroxisomes with UBXD8 localization. 15-20 cells were analyzed in N=3 independent experiments. Scatter plot shows mean and std.dev. ns: not significant, NS: Not significant. ****: P < 0.0001. One-way ANOVA with Dunnett’s multiple comparisons test. Scale bar is 5μM.

**Supplementary Figure 5: Torin1 treatment induces loss of peroxisomes. A.** HeLa wildtype and UBXD8 KO cells were treated with Torin1 (1μM for 16hrs) stained for peroxisomes using catalase. (**B)** Quantification of peroxisomes per cell. 100-150 cells were analyzed in N=3 independent experiments. Violin plot shows median and 95% confidence intervals. **, **** : P < 0.01, 0.0001. Two-way ANOVA with Dunnett’s multiple comparisons test. Scale bar is 5μM.

**Supplementary Figure 6: Depletion of ATG5 rescues peroxisome abundance. A.** HeLa wildtype and UBXD8 KO cells were depleted of ATG5 using siRNA. Cells were stained for peroxisomes using catalase. **B.** Quantification of peroxisome abundance from (A). 100-150 cells were analyzed in N=3 independent experiments. Violin plot shows median and 95% confidence intervals. *, **** : P< 0.05, 0.0001. Two-way ANOVA with Dunnett’s multiple comparisons test. Scale bar is 10 μM. **C.** Immunoblot showing ATG5 depletion.

## Material and Methods

### Cell Culture and Treatments

HeLa-Flp-IN-TREX (HFTs (ThermoFisher Cat# R71407) with introduced Flp-In site (Flp-In™ T-REx™ Core Kit, Cat# K650001; Thermofisher Scientific is a gift from Brian Raught, University of Toronto), COS7 and HEK-293T (ATCC) cells were cultured in Dulbecco’s modified Eagle’s medium, supplemented with 10% fetal bovine serum (FBS) and 100 units/ml penicillin and streptomycin. Cells were maintained in a humidified, 5% CO2 atmosphere at 37 °C. For siRNA transfections, cells were either forward or reverse transfected with 20 nM siRNA using Lipofectamine RNAiMax (Invitrogen) in a 12- or 6-well plates according to the manufacturer’s protocol. After 24 or 48 hours depending on the study, cells were split into 12-well plates for further analysis. 48 or 72 hours post transfection, cells were harvested for immunoblot or fixation for immunofluorescence. For DNA transfections, 0.5 µg HA- and FLAG-tagged wildtype UBXD8 and domain deletions, 0.5 µg HA-/FLAG-tagged UBXD8 UBX domain mutant (FPR to AAA), 0.5 µg HA-/FLAG-tagged UBXD8 UAS domain deletion, 0.5 µg HA-/FLAG-tagged UBXD8 UBA domain mutant (LLQF to AAAA), 0.25 µg GFP-USP30, 0.25 µg RFP-SKL, 0.5 µg GFP-Cherry-PEX26_TM_ constructs were forward transfected into cells seeded in either a 6-well or 12-well plate using Lipofectamine 2000 (Invitrogen). The cells were then harvested 48 or 72 hours post transfection. Cells were lysed in mammalian cell lysis buffer (50 mM Tris-Cl, pH 6.8, 150 mM NaCl, 0.5% Nonidet P-40, HALT Protease inhibitors (Pierce) and 1 mM DTT). Cells were incubated at 4 °C for 10 min and then centrifuged at 19,000 × g for 15 min at 4 °C. The supernatant was collected, and protein concentration was estimated using the DC protein assay kit (Biorad).

### Antibodies and Chemicals

The p97 (10736-1-AP; WB: 1:2000), UBXD8 (16251-1-AP; WB: 1:2000), UBXD2 (21052-1-AP; WB: 1:2000), HRD1 (13473-1-AP; WB: 1:2000), AMFR/GP78 (16675-1-AP; WB: 1:2000), VAPB (14477-1-AP; WB: 1:2000), PEX5 (Eun et al., 2018) (12545-1-AP; WB 1:500), PEX19 (14713-1-AP; WB 1:1000), MLYCD (15265-1-AP; WB 1:2000), PECR (14901-1-AP; WB 1:1000), DECR (25855-1-AP; WB 1:1000), PMP70/ABCD3 (66697-1-Ig; WB 1:1000; IF 1:400), PEX3 (10946-1-AP; WB 1:1000), ACBD5 (21080-1-AP; WB 1:1000), GFP(66002-1-AP; WB: 1:2000) and ATG5 (10181-2-AP; WB 1:1000) antibodies were from Proteintech Inc. The pan-ubiquitin (P4D1; sc8017; WB: 1:2000), c-Myc (9E10; sc40; WB: 1:2000), β-Actin (AC-15; sc69879; WB: 1:2000), and GAPDH (O411; sc47724; WB: 1:2000) antibodies were obtained from Santa Cruz Biotechnologies. LC3B (D11; 3868S; WB: 1:1000), Catalase (12980; WB 1:1000; IF 1:800), and BiP (C50B12; 3177T; WB: 1:2000) were from Cell Signaling Technologies. p97 (A300-589A; WB: 1:2000) was from Bethyl Laboratories. The following antibodies anti-HA (16B12; MMS-101P, Covance; WB: 1:2500), anti-FLAG (M2; F3165 Sigma Aldrich; WB: 1:5000), were used for immunoblotting. HRP conjugated anti-rabbit (W401B; WB: 1:10,000) and anti-mouse (W402B; WB: 1:10,000) secondary antibodies were from Promega. Goat anti-Mouse IgG (H + L) Cross-Adsorbed Secondary Antibody, Alexa Fluor™ 568 (Catalog # A-11004; IF: 1:10,000), and Goat anti-Mouse IgG (H + L) Cross-Adsorbed Secondary Antibody, Alexa Fluor™ 488 (Catalog # A-11001; IF: 1:10,000) were purchased from Thermofisher Scientific. CB-5083 was from Selleckchem. siRNAs were purchased from Ambion (Thermo Fisher Scientific): UBXD8-0 (s23260), UBXD8-9 (s23259). HRD1-3 (D-007090-03), and HRD1-4 (D-007090-04) were purchased from GE Dharmacon. siControl (SIC001) was from Millipore Sigma. p97 siRNAs (2-HSS111263 and 3-HSS111264), UBXD8-C-HA/FLAG construct was previously published (Raman, Havens, Walter, & Harper, 2011). The UBXD8 rescue constructs, including UBA* (^17^LLQF^20^ mutated to ^17^AAAA^20^), ΔUAS (deleted amino acids between 122-277), and UBX* (^407^FPR^409^ mutated to ^407^AAA^409^), were cloned using overlap PCR followed by Gibson assembly (NEB) cloning into pHAGE-C-HA/FLAG. Torin1 (502050475) and Clofibrate (08-241-G) are from Fisher Scientific. Cycloheximide (97064-724) is from VWR International.

### Immunofluorescence and Microscopy

HFT and HEK293T cells were plated on # 1.5 glass coverslips in a 12-well plate. Following indicated treatments, cells were fixed in 4 % paraformaldehyde (PFA) (15710-S Electron Microscopy Sciences) diluted in PBS for 15 minutes at room temperature. Next, cells were washed in PBS and permeabilized in ice-cold 100% methanol at − 20 °C for 10 min. Cells were then washed three times in PBS and incubated in blocking buffer (1 % BSA, 0.3 % Triton-X100 for 1 hour at room temperature. Primary antibodies were diluted to the indicated concentrations in blocking buffer and coverslips were incubated overnight at 4 °C in a humidified chamber. Coverslips were then washed three times in PBS and incubated in secondary antibodies diluted to the indicated concentrations in blocking buffer for 1.5 hour at room temperature. The secondary antibody solution was replaced with Hoechst diluted in blocking buffer and incubated for 5 minutes at room temperature.

Coverslips were washed three times with PBS and mounted to slides with ProLong Gold antifade mounting media (P36930 Invitrogen). All images were collected using Zeiss LSM800 confocal microscope equipped with Airyscan. Images were taken at 63 X (with oil) magnification. The indicated fluorophores were excited with a 405, 488, or 594 nm laser line.

### Image analysis

Images were analyzed using FIJI (https://imagej.net/fiji). Peroxisome number per cell and size was measured using an automated image analysis script Aggrecount which allowed for segmentation and single cell resolution (Klickstein et al., 2020). ImageJ JACOP plugin was used for colocalization analysis. Total number of peroxisomes in each cell were counted using ImageJ “analyze particle” tool. Peroxisomes (red) colocalization with UBDX8 (green) were determined from merging images from red and green channels. The color threshold tool was used to select yellow puncta (Hue value 25-60) that indicates colocalization and quantified using ImageJ “analyze particle” tool.

For pexophagy flux reporter assays, the background was subtracted, and ROIs were generated based on the mCherry signal. ROIs were transferred to the GFP channel and the GFP intensity was measured. Images in each replicate were carefully examined and GFP threshold intensity was empirically determined for all images in a given replicate. The number of puncta with GFP above the pre-determine intensity was calculated for the ratio.

### TMT Proteomics and Lipidomics

The proteomic and lipidomic data were previously published in (Ganji et al., 2023). Please refer to the Supplementary information in that manuscript for the individual datasets. Raw data is available via the ProteomeXchange Consortium via the PRIDE (Perez-Riverol et al., 2022) partner repository with the dataset identifier PXD-39061. The mass spectrometry lipidomics data is available at the NIH Common Fund’s National Metabolomics Data Repository (NMDR) website, the Metabolomics Workbench, (https://www.metabolomicsworkbench.org), where it has been assigned Project ID PR001559 [10.21228/M85X3W] with StudyIDs ST002421.

### Gene ontology (GO) functional enrichment analyses of proteomics data

The differentially expressed proteins were further annotated and GO functional enrichment analysis was performed using Metascape online tool (http://metascape.org) (Zhou et al., 2019).

The GO cluster network and protei-protein interaction network generated by Metascape. Other proteomic data visualizations were performed using the RStudio software (v1.4.1103).

### Quantitative PCR

For all real-time PCR experiments, total RNA was isolated using the Quick-RNA Miniprep Kit (Zymo Research cat. no. R1055). The purified RNA was quantified by NanoDrop and 1 µg of RNA for each sample was used to generate cDNA using the iScript cDNA synthesis kit (Biorad cat. no.1708890). Real-time PCR was performed with PowerUp SYBR Green Master Mix (Applied Biosystems cat. no. A25741) on an Applied Biosystems StepOnePlus real-time PCR system. Data analyses utilized the 2 ^-ΔΔCt^ method and GAPDH was used as a housekeeping gene to normalize transcript expression across samples. The XBP1s primers were previously published (van Schadewijk, van’t Wout, Stolk, & Hiemstra, 2012) as well as all peroxisomal primers (Bagattin et al., 2010). Primers are as follows:

ACOX1 F(CCATTCAAGCTGTCTTAAGGAGTT), R(CTGAGGCTCTGTCATGATGC).

ACOX2: F(CAAATTGTCGGCCTCCTGTA), R(GAGATCTCTGTGGCGTGGAG).

PBFE: F(AAGAAGGACTACAGAAAGCTGTA, R(CCCAGTGTAAGGCCAAATGT).

DBP: F(GTGGCTTGTTTGAGGTTGGA), R(CCTCAGGAGTCATTGGGTGA).

PTHIO: F(TACTTCGCGCTTGATGGAGA), R(TCTCCCGTGAAATGCCAAAC).

Pex3: F(TTCTTTTGCGGGTCCAGTTA), R(ACATCTGGGGGAGCAAGAAT).

Pex7: F(TCTGGCTCATGGGATCAAAC), R(GGATGTGGGGAGACCAGATT).

Pex12: F(AAGCTCTGGAGCACAAACCA), R(ACACCCCCAACAGCTTTCTT).

Pex13: F(CCGGGCTGGTGATATGCT), R(GTATAAGTCCTGTTGTTTGGCCATC).

Pex16: F(CGAGCTGTCAGAGCTGGTGTACT), R(ACAGCGACACAGGCAACTTTT).

Pex19: F(CTCTCAGAGGCTGCAGGGAG), R(GTGGCATTTTTGGCTAATCCA).

Fis1: F(AAAGGGAGCAAGGAGGAACAG), R(AACCCGCGGACGTACTTTAAG).

DLP1: F(TCGTCGTAGTGGGAACGCA), R(TCTCCGGGTGACAATTCCAG).

XBP1s: F(TGCTGAGTCCGCAGCAGGTG), R(GCTGGCAGGCTCTGGGGAAG).

Total XBP1: F(AAACAGAGTAGCTCAGACTGC), R(TCCTTCTGGGTAGACCTCTGGGAG).

### Catalase activity assay

Catalase activity was determined using the Catalase Assay Kit (ab83464, Abcam) as per manufacturer’s instructions. Briefly, cells were lysed and protein concentration of the cell lysate was determined. Catalase decomposes H_2_O_2_ to water and oxygen. The assay uses the unconverted H_2_O_2_ and reacts with OxiRed probe to produce a product that can be measured at 570 nm. The catalase activity present in the sample is inversely proportional to the signal obtained. The kit can detect as little as 1 µU of catalase activity.

### Statistics and reproducibility

For all experiments, n ≥ 3 or more biological replicates for each condition examined. Fold changes, SEM, SD, and statistical analyses were performed using GraphPad Prism version 9.4.1 for Windows (GraphPad Software). Statistical tests and N values are mentioned in the figure legends.

## Acknowledgements

We thank Xiaosheng Yang and Jacob Liebovitz for critical reading of the manuscript. We thank Rakesh Ganji for producing the TMT proteomics data set and the UBXD8 KO cell lines. We thank James Olzmann for the GP78 KO cell line, and Peter Kim for the flux reporter construct. We are grateful to Vibha Ramu and Ly Nguyen for help with immunoblots and image analysis. This work is supported by the NIH grants R01 GM127557 and R21 NS123631 to M.R., R35 GM147189 to A.J and NRSA F31 GM148057-01 to I.D.M.

## Respective Contributions

I.D.M and M.R conceived the studies. I.D.M performed all studies and data analysis with assistance from M.R. Imaging and analysis of UBXD8 localization to peroxisomes was performed by S.A with assistance from A.J. I.D.M and M.R wrote the manuscript.

## Request for reagents

Please contact the corresponding author, M.R for reagent requests.

## Data availability

Raw data is available via the ProteomeXchange Consortium via the PRIDE (Perez-Riverol et al., 2022) partner repository with the dataset identifier PXD-39061. The mass spectrometry lipidomics data is available at the NIH Common Fund’s National Metabolomics Data Repository (NMDR) website, the Metabolomics Workbench, (https://www.metabolomicsworkbench.org), where it has been assigned Project ID PR001559 [10.21228/M85X3W] with StudyIDs ST002421.

