## Supplementary figures and images for "The p97-UBXD8 complex maintains peroxisome abundance by suppressing pexophagy"

### Supplemental Figure 1

Supplementary Figure 1.

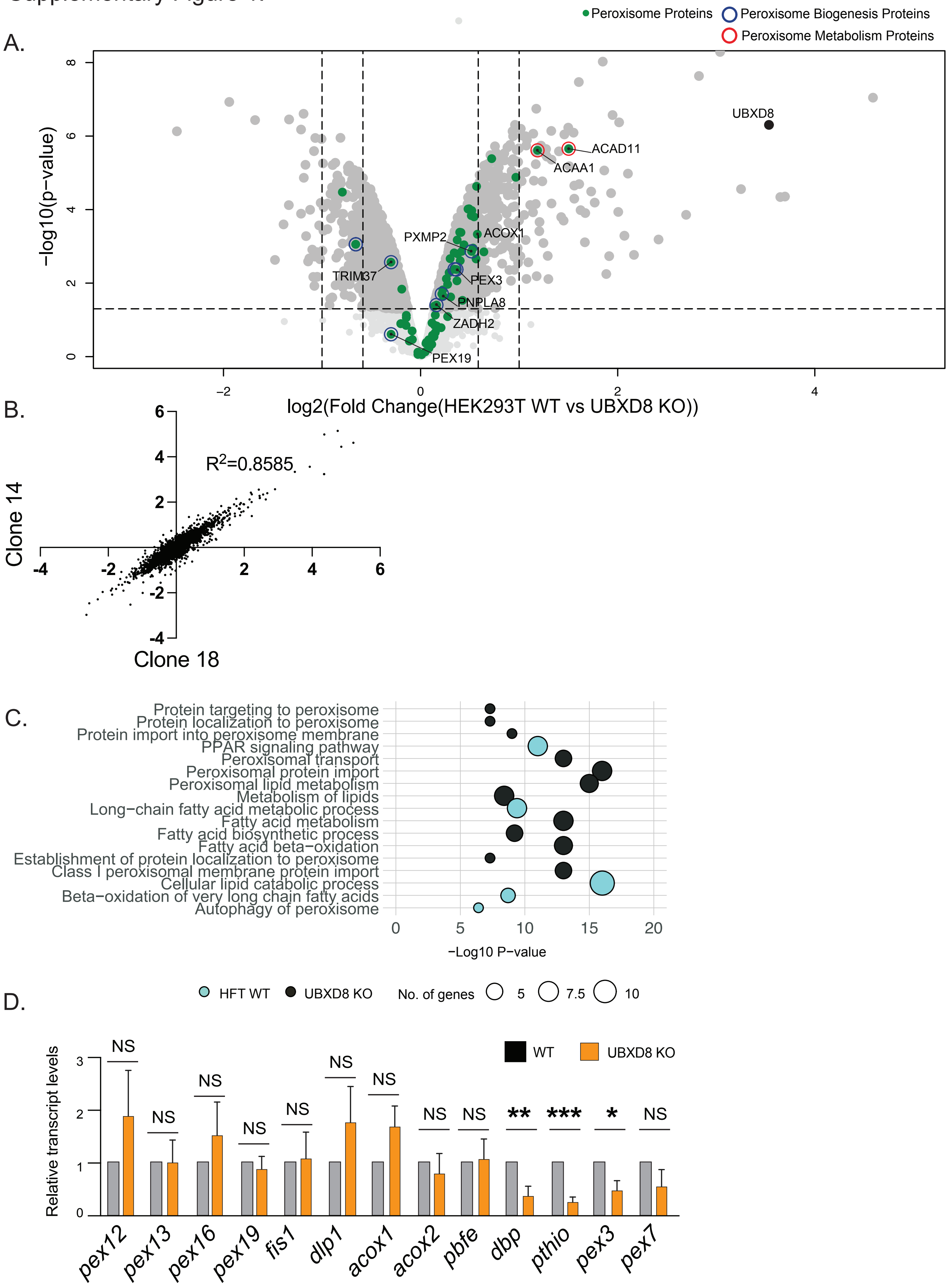

### Supplemental Figure 2

Supplementary Figure 2.

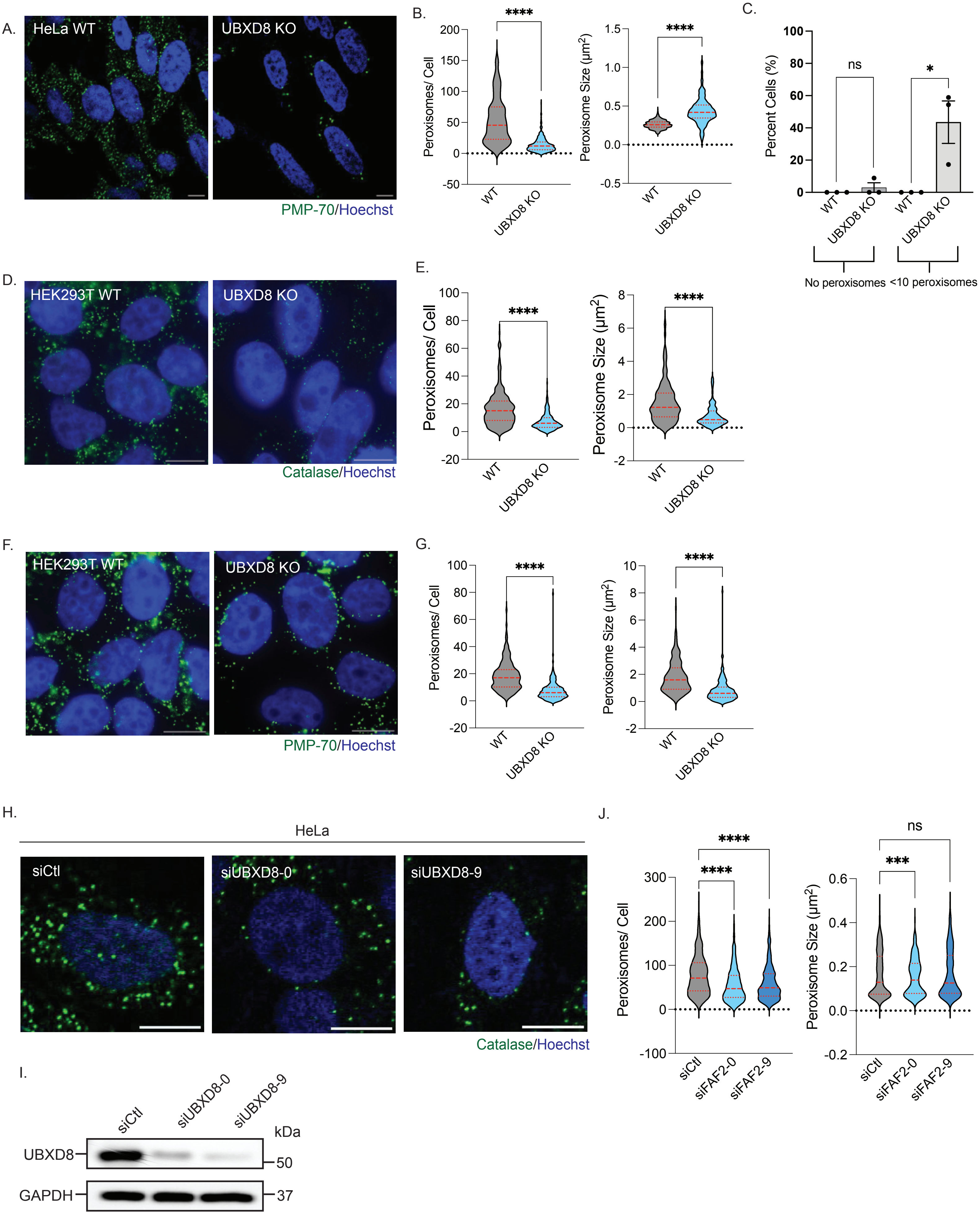

### Supplemental Figure 3

Supplementary Figure 3.

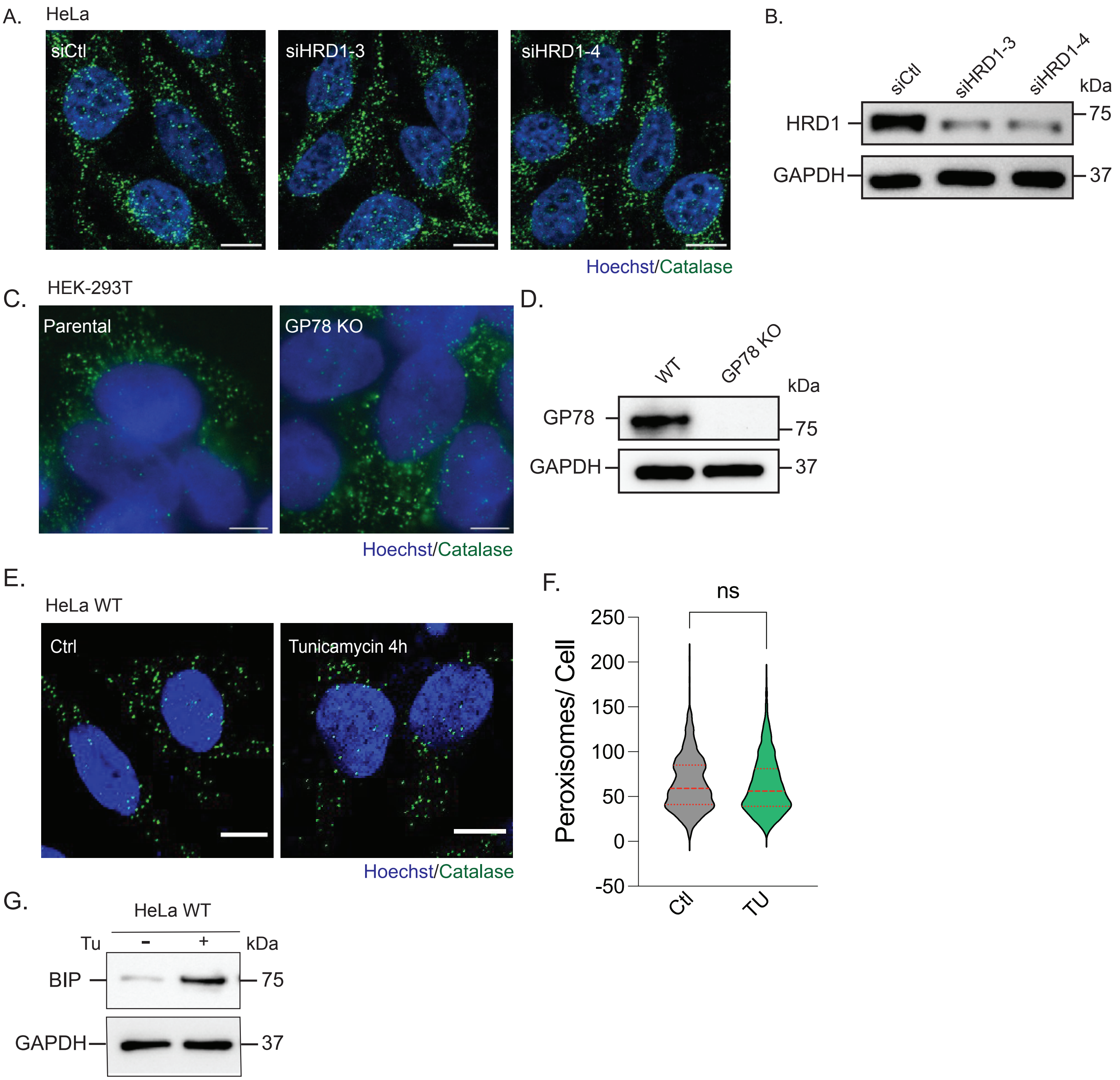

### Supplemental Figure 4

Supplementary Figure 4

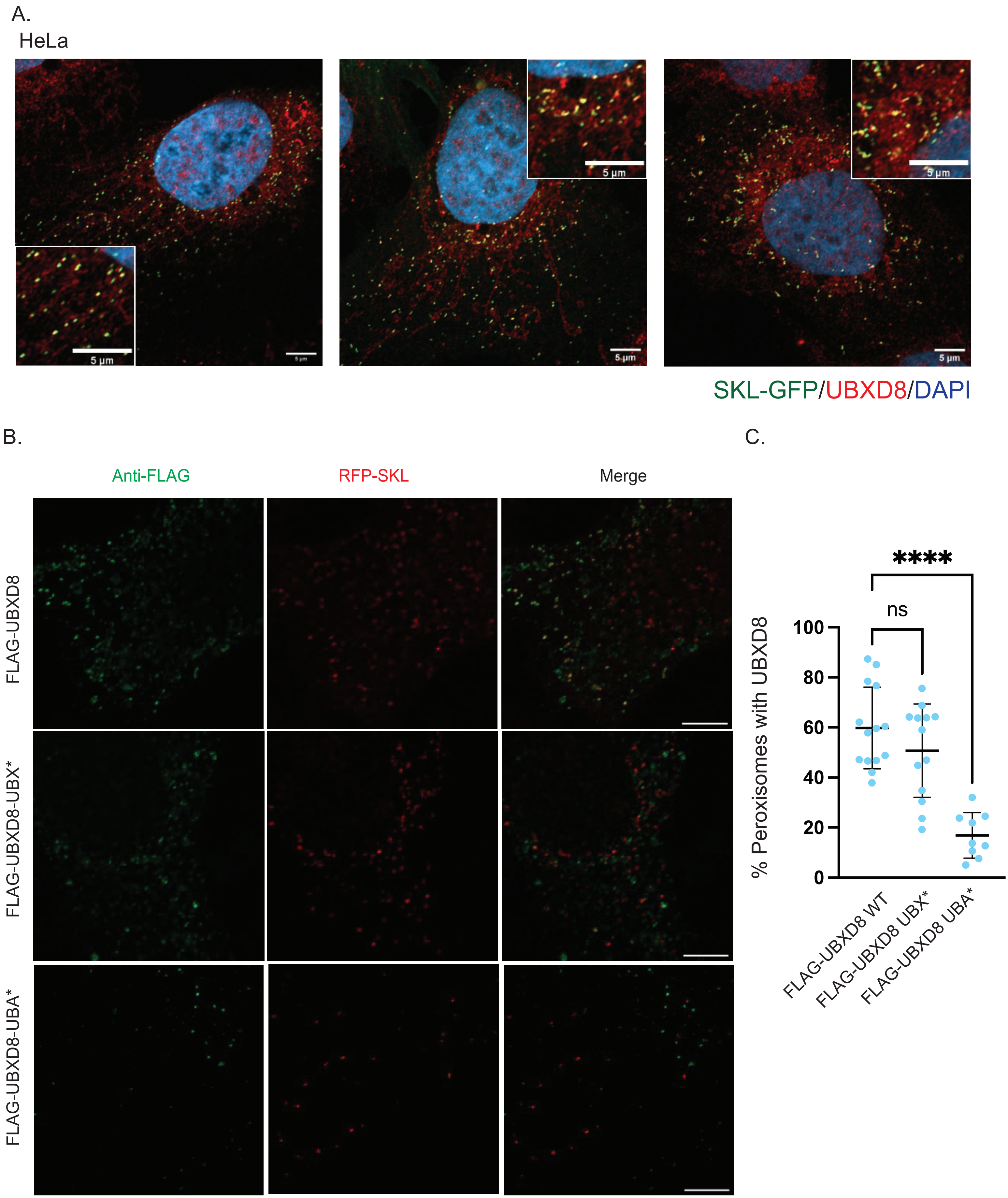

### Supplemental Figure 5

Supplementary Figure 5

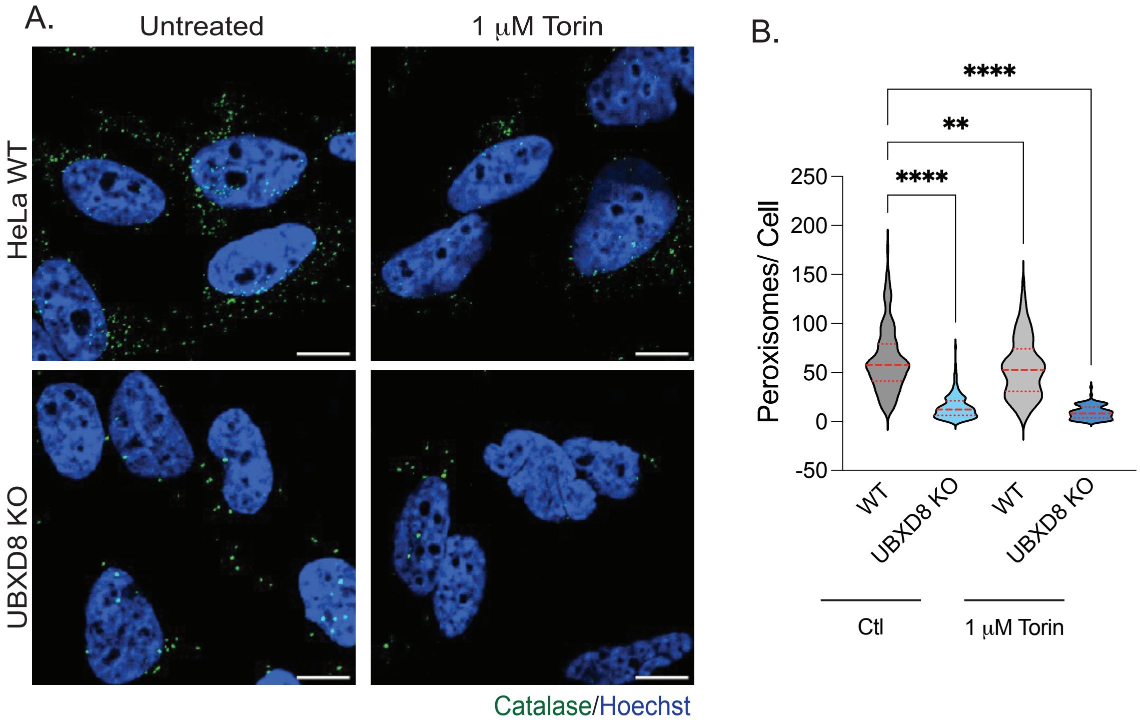

### Supplemental Figure 6

Supplementary Figure 6.

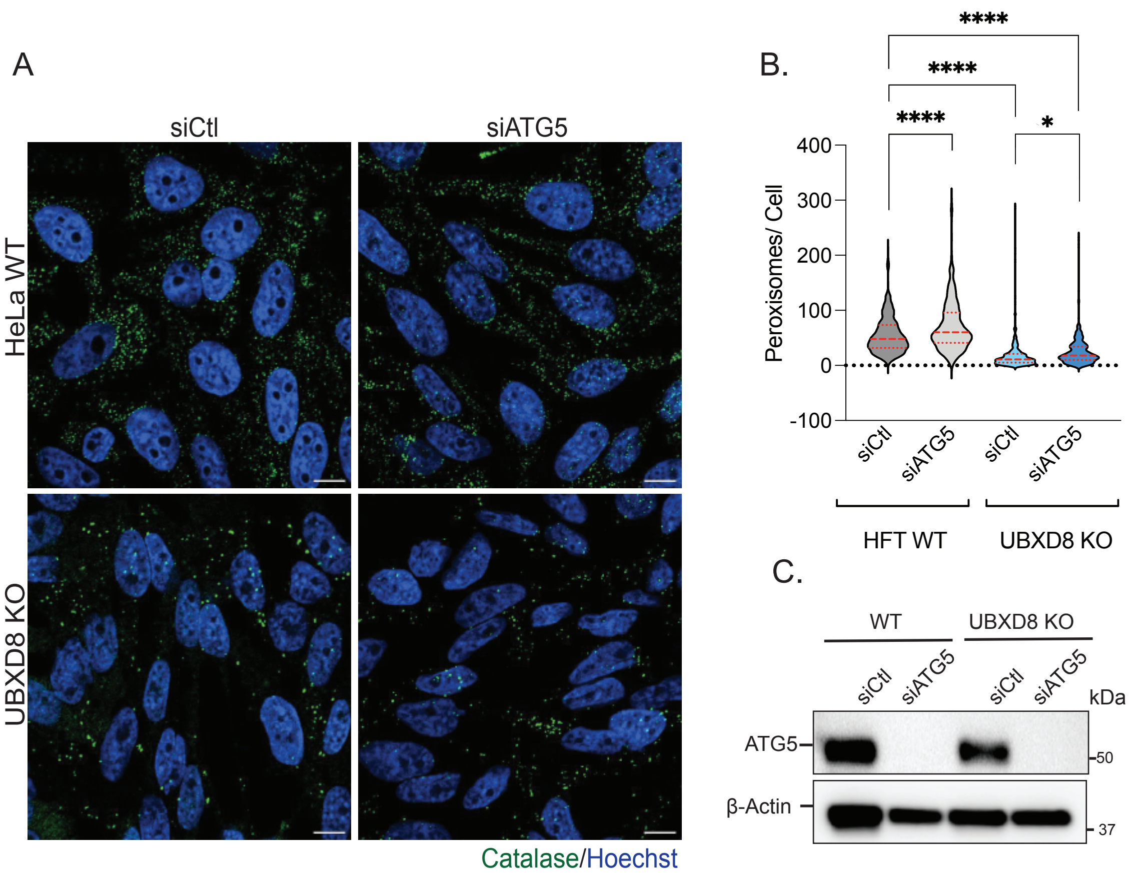
